# Triplex Domain Finder: Detection of Triple Helix Binding Domains in Long Non-Coding RNAs

**DOI:** 10.1101/020297

**Authors:** Sonja Hänzelmann, Chao-Chung Kuo, Marie Kalwa, Wolfgang Wagner, Ivan G. Costa

## Abstract

Long (>200vbps) non-coding RNAs (lncRNA) can act as a scaffold promoting the interaction of several proteins, RNA and DNA. Some lncRNAs interact with the DNA via a triple helix formation. Triple helices are formed by a single stranded RNA/DNA molecule, which binds to the major groove of a double helix following a canonical code. Recently, sequence analysis methods have been proposed to detect triple helices for a given RNA and DNA sequences. We propose the Triplex Domain Finder (TDF) to detect DNA binding domains in RNA molecules. For a candidate lncRNA and potential target DNA regions, i.e. promoter of genes differentially regulated after the knockdown of the lncRNA, TDF evaluates whether particular RNA regions are likely to form DNA binding domains (DBD). Moreover, the DNA binding sites from the predicted DBDs are used to indicate potential target DNA regions, i.e. genes with high binding site coverage in their promoter. The command line tool provides results on a user friendly and graphical html interface. A case study on FENDRR, an lncRNA known to form triple helices, demonstrates that TDF is able to recover both previously discovered DBDs and DNA binding sites. Source code, tutorial and case studies are available at www.regulatory-genomics.org/tdf.

## 1 Introduction

Long non-coding RNAs (lncRNAs) represent a novel type of RNA molecule with two distinct characteristics: they are not translated into proteins (non-coding) and can have a size up to thousands of nucleotides. These molecules have complex biological functions, such as their ability to interact with any other molecule in the cell (DNA, RNA and proteins). Due to their length they enhance simultaneous interactions of many molecules. The few well-studied lncRNAs have been shown to participate in important biological processes, such as cell development and diseases by changing the packing of chromosomes and the activation of closed genes (Mercer and Mattick, 2013).

Two recent techniques measure the interaction of particular lncRNAs with DNA: Chromatin Isolation by RNA purification followed by sequencing (ChiRP-Seq) (Chu et al., 2011b) and capture hybridization analysis of RNA targets (CHART-Seq) (Simon et al., 2011). For example, ChiRP-Seq indicates that the lncRNA HOTAIR interacts with more than 900 genomic regions in a human cancer cell line and that these interactions occur close to sites with EZH2, SUZ12 and H3K27me3 occupancy (Chu et al., 2011b). An alternative is the use of loss of function assays with shRNAs, which have demonstrated the importance of FENDRR in mouse mesoderm differentiation (Grote et al., 2013). However, understanding the exact molecular mechanisms of lncRNAs requires extensive and laborious work. An alternative is the use of computational tools to predict particular molecular interactions, such as potential triple helix binding sites of lncRNAs to the DNA.

A triple helix is formed between a single stranded RNA molecule and the major groove of a double helix DNA via Hoogsten hydrogen bonds (or double helix RNA) (Felsenfeld et al., 1957). The triple helix can be formed in two configurations: the purine rich DNA strand and the RNA are in the same orientation (5 to 3, parallel) or in anti-parallel (5 to 3 and 3 to 5) orientations. Three combinations of bases promote stable triple helices (RNA-DNA:DNA): in the parallel configuration the stable combination of the bases are: T-A:T; C-G:C, G-G:C and in the anti-parallel configuration: A-A:T, G-G:C and T:A:T.

Recently, Buske et al. (2012) proposed a sequence analysis method (Triplexator) to predict short (<30 bps) DNA binding sites in the genome for a given RNA using the previously described canonical rules. The algorithm efficiently detects triple helix complexes in large DNA regions. Latter, a methodology that detects longer triple helix binding sites and DNA binding domains in single DNA regions has been proposed (He et al., 2015). However, none of the methods is able to statistically evaluate the triple helix forming potential of lncRNAs in multiple DNA regions as provided by genome-wide functional studies previously described.

To address this problem, we propose a statistical approach, Triplex Domain Finder (TDF). Our tool detects enriched DNA binding domains (DBDs) on the RNA and ranks DNA target regions by DNA binding site (DBS) statistics.

## 2 Method

### 2.1 Overview

Triplex Domain Finder (TDF) starts by finding DNA binding sites (DBS) and RNA binding sites (RBS) of a lncRNA in DNA regions. One set of regions comprises the potential lncRNA targets (target regions), e.g. promoters of genes differentially expressed after FENDRR knockdown, and a second set defined as non-target regions, e.g. promoters of genes not differentially expressed. Next, we list all candidate DNA binding domains (DBDs) by finding regions in the RNA with overlapping RBS (see Figure 1). We only consider candidate DBDs with more than k DBS in the target regions (k = 20 as default). Finally, we test whether the number of target regions with at least one DBS is higher than the number of non-target regions with at least one DBS for a given DBD.

**Figure 1:**
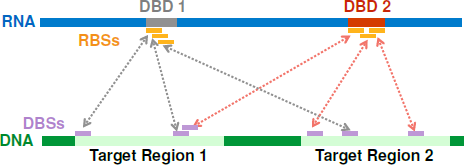
Schematic of binding site nomenclature used in TDF for an RNA and a set of DNA target regions. Each triple helix is formed by one RNA binding site (RBS) and one DNA binding site (DBS). The DNA binding domains (DBDs) are RNA regions with overlapping RBSs. In the example, we have two candidate DBDs forming a potential triple helix with several DBSs in two DNA target regions.

### 2.2 Definitions

The fundamental operations are based on a **genomic region**, which is a linear interval with specific positions in the genome, defined as follows:

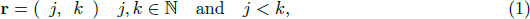

where **r** denotes a linear interval in the genome with *j* and *k* as the start and end excluding positions. A **genomic region set** is defined as a collection of *n* genomic regions.

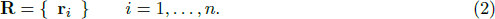

For a pair of regions *r* = (*j*, *k*) and e = (*l*, *m*), we can define an overlap as

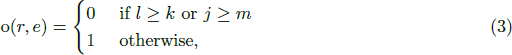

and merge operation as

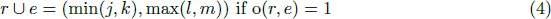

A triple helix binding site is defined as a tuple of an RBS (RNA Binding Site) and a DBS (DNA Binding Site)

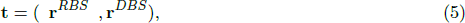

where **r**^*RBS*^ is the binding site location in the RNA and **r***^DBS^* is the binding site location in the DNA. The collection of all binding events is defined as

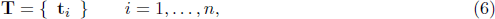

where *n* is the total number of the binding events. Similarly, we define all RBSs and DBSs as **R**^*RBS*^ and **R**^*DBS*^

### 2.3 Promoter Test

The promoter test evaluates the probability of a triple helix complex forming in the promoter regions of candidate genes (e.g. genes differentially expressed in a particular functional study). The test compares the binding events of the lncRNA in the promoters of candidate genes (target regions) with the binding events in the remaining promoters of the genome (non-target regions).

Given the set of target regions **R**^*tr*^, the non-target regions **R**^*ntr*^ and a lncRNA, we use Triplexator (Buske et al., 2012) to predict the triple helix binding sites in target DNA regions **T**^*t*^ and non-target regions **T**^*nt*^. Next, we list all potential DNA binding domains by finding regions in the lncRNA with overlapping RBSs associated to the target regions (see Figure 1 for a visual representation).

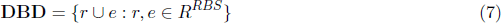

TDF ignores DBDs having a low number of DBSs (< 20 as default). The candidate DBDs are mutually exclusive. Next, we define a function to enumerate all DBSs from a set of triple helix predictions *T* associated with a DBD *d.*

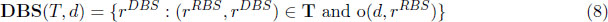

Then, we count the number of target regions with at least one DBS for a given DBD *d.*

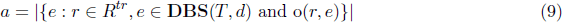

and non-target regions with at least one DBS for a given DBD *d.*

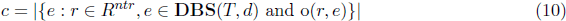

We then define a two by two contingency table representing the number of targets (non-targets) regions with at at least one (or no) DBS for a given DBD.

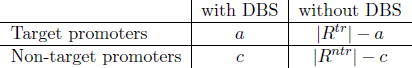

Finally, the Fisher’s Exact test is performed on the above 2×2 contingency table for each DBD. TDF reports both the corrected *p*-value (Benjamini and Hochberg, 1995) and odds-ratio statistics.

#### 2.3.1 Genomic Region Test

The genomic region test evaluates the triple helix binding potential of a lncRNA with predefined DNA regions (target regions). Target regions can be obtained using genome-wide assays that measure the interaction of lncRNAs with the DNA (ChiRP-Seq and CHART-Seq).

Due to a lack of non-target regions, we generate *L* non-target regions by random selection of genomic regions with the same size/length as the target region. We use an empirical statistical test to evaluate whether the number of target regions with at least one DBS is larger than the number of “random” non-target regions with a minimum of one DBS for a given DBD.

First, we generate *L* non-target regions by randomly selecting DNA positions of the same size as the target regions. Then, we run the Triplexator on the regions and obtain the predictions *T^tr^* and 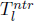 for *l* = 1,…, *L.*

Next, we evaluate all potential DBSs from the target regions as described in Equation 8. Then, we estimate the number of target regions (*a*) with at least one DBS for a given DBD (Equation 9). Similarly, we obtain a distribution **c** = {*c*,.., *c_l_*,.., *c_L_*} where *c_l_* is the number of non-target regions with at least one DBS per DBD of the *l*th non-target region. Finally, we compute an empirical *p*-value by counting the number of values higher than *a* that are found in **c**.

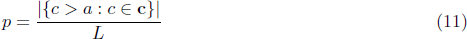

Finally, we apply the false discovery rate (FDR) (Benjamini and Hochberg, 1995) as multiple test correction method to the *p-*values.

#### 2.3.2 Genomic Annotation

The promoter regions are obtained from gene annotations for mouse (mm9) and for human (hg19). TDF accepts either gene symbols or ENSEMBL IDs as input lists for the promoter test. Mapping of genes symbols to ENSEMBL is made based on annotations provided by HGNC for human (http://www.genenames.org/) and by MGI for mouse (http://www.informatics.jax.org).

#### 2.3.3 Ranking of regions and promoters

TDF also provides simple approaches to rank target regions and promoters. Two statistics are based on DBS: the number of DBSs associated with the region and the percentage of the base pairs covered by at least one DBS. TDF also allows the inclusion of experimental evidence, e.g. gene expression fold change or scores of ChiRP-Seq peaks, as an external criteria for ranking. TDF provides a combined statistic using the sum of ranks approach. Moreover, TDF produces as result an HTML interface allowing the user to select the criteria for ranking candidate target regions. For the promoter test, TDF also shows the genes associated with the promoter. For the genomic region test, for each of the genomic regions the closest upstream and downstream genes in the neighborhood of at least 50kbs of the region.

## 3 Results

### 3.1 Fendrr

Grote et al. (2013) identified the lncRNA FENDRR and studied its involvement in mouse development using knock down experiments. They demonstrate that FENDRR plays a role in ES cell differentiation via gene chromatin modifications. They also show that FENDRR forms a triple helix in the promoters of *Foxf1* and *Pitx2*.

To define genes potentially targeted by FENDRR, we obtained the FPKM (fragments per kilo-base per million fragments mapped) table provided in Gene Expression Omnibus (GSE43078). We calculated the log2 fold change between FENDRR siRNA and Control. Genes with a 2-fold change are defined as differentially expressed (1507 genes). We also included genes analyzed by Grote et al. (2013), which were not present in our gene list.

We used the promoter test to detect potential triple helix binding sites of FENDRR in the upstream regions of each gene (1kb). Triple helix binding sites had at least 15 bps with up to 20% of error. As shown in Figure 2 and Table 1, we find 4 significantly enriched DNA binding domains. The DBS with the smallest p-value (2.0e-12; 1502-1565) matches the DBD experimentally verified in Grote et al. (2013). Moreover, TDF ranking criteria indicate PITX2 as the 4th gene with the highest number of DBSs and FOXF1 as also the 4th taking into account the sum of ranks (Table 2).

**Figure 2:**
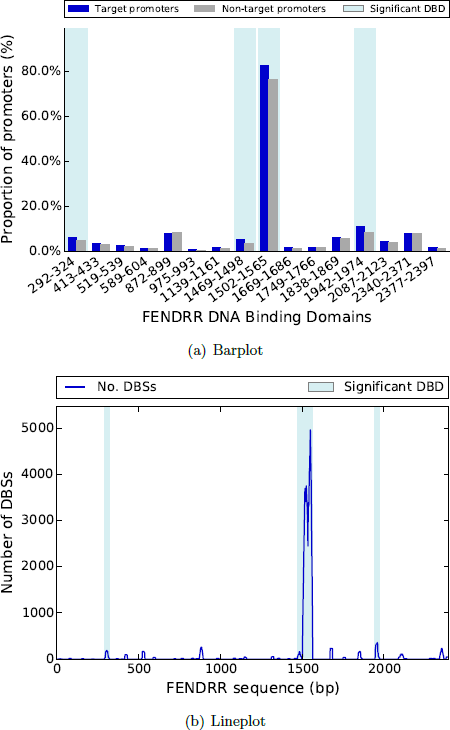
a) The bar plot shows the proportion of target and non-target regions with at least one DBS (y-axis) associated to a given DBD (x-axis) for FENDRR. b) depicts the number of promoters targeted by a DBS (y-axis) in the location of the associated RBS on the RNA (x-axis). Four DBDs have a significantly higher number of DBSs in targets than non-target regions (highlighted in blue).

**Table 1:** Candidate DBDs from FENDRR ranked by *p*-values.

|  | Target Promoters |  | Non-Target Promoters |  | Statistics |  |
| --- | --- | --- | --- | --- | --- | --- |
| DBD Position | DBS | no DBS | DBS | no DBS | Odds ratio | *p*-value |
| 1502-1565 | 1919 | 400 | 42635 | 13189 | 1.48 | 2.0e-12 |
| 1942-1974 | 262 | 2057 | 4804 | 51020 | 1.35 | 6.6e-05 |
| 1469-1498 | 126 | 2193 | 2181 | 53643 | 1.41 | 0.0014 |
| 292-324 | 145 | 2174 | 2839 | 52985 | 1.24 | 0.0339 |
| 2377-2397 | 46 | 2273 | 894 | 54930 | 1.24 | 0.29 |
| 519-539 | 70 | 2249 | 1443 | 54381 | 1.17 | 0.30 |
| 975-993 | 21 | 2298 | 398 | 55426 | 1.27 | 0.39 |
| 413-433 | 81 | 2238 | 1814 | 54010 | 1.08 | 0.40 |
| 589-604 | 37 | 2282 | 778 | 55046 | 1.15 | 0.40 |
| 1669-1686 | 40 | 2279 | 872 | 54952 | 1.11 | 0.40 |
| 1838-1869 | 148 | 2171 | 3406 | 52418 | 1.05 | 0.40 |
| 2087-2123 | 103 | 2216 | 2305 | 53519 | 1.08 | 0.40 |
| 1139-1161 | 40 | 2279 | 889 | 54935 | 1.08 | 0.41 |
| 1749-1766 | 46 | 2273 | 1090 | 54734 | 1.02 | 0.51 |
| 2340-2371 | 189 | 2130 | 4501 | 51323 | 1.01 | 0.51 |
| 872-899 | 190 | 2129 | 4764 | 51060 | 0.96 | 0.73 |

**Table 2:**
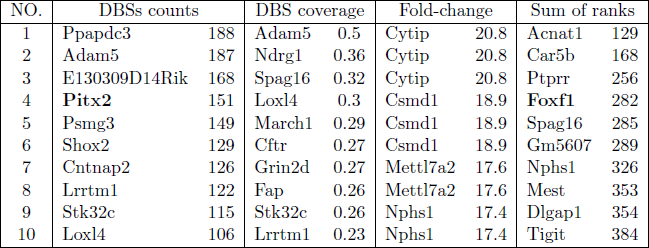
Top 10 gene promoters from different ranking strategies.

### 3.2 Terc

We obtained regions from TERC ChiRP-Seq on HeLa cells (Chu et al., 2011a) from the Gene Expression Omnibus (GSE31332). We used the genomic region test to estimate whether TERC forms triple helices within the 2198 target regions found with ChiRP-seq. Triple helix binding sites have at least 15 base pairs and maximum error of 20% (default). We used L = 10,000 random nontarget regions. As shown in Figure 3 and Table 3, TDF finds 4 significant DNA binding domains within TERC.

**Table 3:** Candidate TERC DBDs ranked by *p*-values.

|  | Target Regions | Non-Target Regions |  |  |
| --- | --- | --- | --- | --- |
| DBDs position | DBS | DBS (mean) | s.d. | $p$-value |
| 0-34 | 471 | 156.7 | 11.7 | 0 |
| 189-205 | 61 | 28.3 | 5.39 | 0 |
| 332-348 | 105 | 24.1 | 4.86 | 0 |
| 360-387 | 169 | 104.3 | 10.0 | 0 |
| 161-178 | 33 | 65.3 | 8.15 | 1.00 |

**Figure 3:**
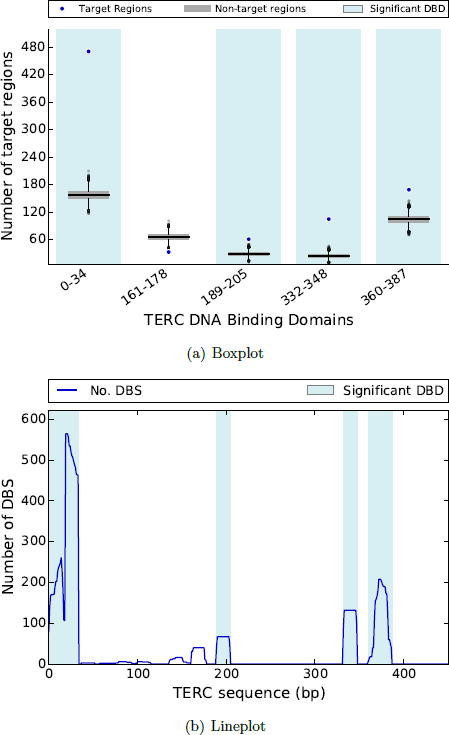
a) The box plot shows the number of target and non-target regions with at least one DBS (y-axis) associated for a given DBD (x-axis) for TERC. Four DBDs have significantly higher number of DBDs in targets than non-target regions (highlighted in blue), b)Number of promoters targeted by DBS (y-axis) vs. the location of the associated RBS on the RNA (x-axis). Significant DBDs are indicated in light blue.

### 3.3 MEG3

We obtained the binding regions of MEG3 in human breast cancer cell line (BT-549) from (Mondal et al., 2015). A chromatin oligo affinity precipitation (ChOP) method followed by sequencing revealed 532 binding sites of Meg3. We applied them to genomic region test of TDF to evaluate the triple helix binding affinity of MEG3 in these regions. As shown in Figure 4 and Table 4, TDF finds seven significant DNA binding domains within MEG3.The DBD with highest number of target DNA regions corresponts to the binding domain, which was show to binding to an enhancer of TGFBR1 via triple helices (Mondal et al., 2015). The enhancer region is however only the top

**Table 4:** Candidate MEG3 DBDs ranked by number of DBSs

| DBDs position | Target Regions |  | Random Regions |  | *p*-value |
| --- | --- | --- | --- | --- | --- |
|  | DBS | no DBS | DBS (average) | s.d. |  |
| 13-41 | 155 | 377 | 15.8 | 3.87 | 0 |
| 961-1008 | 138 | 394 | 28.6 | 5.27 | 0 |
| 839-890 | 99 | 433 | 28.7 | 5.26 | 0 |
| 1458-1498 | 92 | 440 | 4.41 | 2.10 | 0 |
| 693-725 | 78 | 454 | 27.8 | 5.07 | 0 |
| 442-458 | 68 | 464 | 8.57 | 2.87 | 0 |
| 1159-1190 | 68 | 464 | 15.6 | 3.91 | 0 |

**Figure 4.**
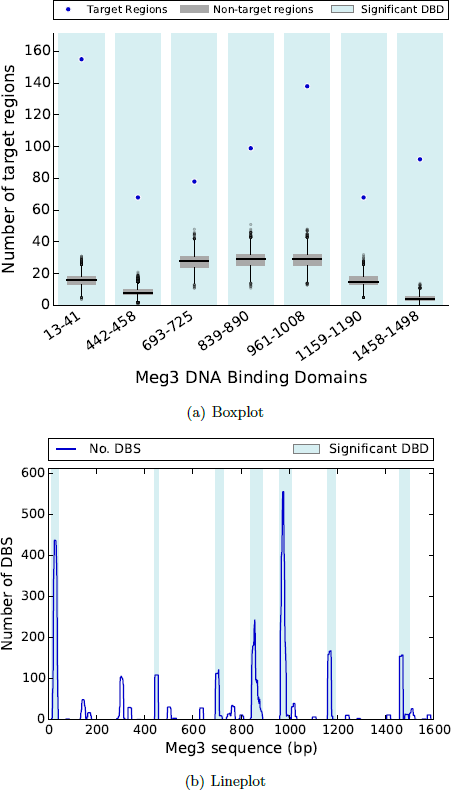
a) The box plot shows the number of target and non-target (random) regions with at least one DBS (y-axis) associated for a given DBD (x-axis) for MEG3. All seven DBDs have significantly higher number of DBDs in targets than the random regions (highlighted in blue), b) Number of target regions targeted by DBS (y-axis) vs. the location of the associated RBS on the RNA (x-axis). Significant DBDs are indicated in light blue.

## 4 Discussion

Long non-coding RNAs have the unique capacity of interacting with several DNA, RNA or proteins simutaneously (Guttman and Rinn, 2012; Johnson and Guig´o, 2014). In this context, the computational characterization of such interaction domains is an very important step for understanding lncRNA function. We describe here the first method to detect DNA interactions domains formed via triple helices by statistical analysis of target DNA regions originated from high throughput functional experiments. Our method is not only able to detect validated DNA binding domain of FENDRR and MEG3 and its targets genes, as it shows statistical evidence that several lncRNAs have triple helix forming domains.

## References

Yoav Benjamini and Yosef Hochberg. Controlling the false discovery rate: a practical and powerful approach to multiple testing. Journal of the Royal Statistical Society. Series B (Methodological), pages 289–300, 1995. ISSN 00359246. URL http://www.jstor.org/stable/2346101.

Fabian A. Buske, Denis C. Bauer, John S. Mattick, and Timothy L. Bailey. Triplexator: detecting nucleic acid triple helices in genomic and transcriptomic data. Genome research, 22(7):1372–1381, 2012. ISSN 10889051. URL http://genome.cshlp.org/content/22/7/1372.short.

Ci Chu, Kun Qu, Franklin L. Zhong, Steven E. Artandi, and Howard Y. Chang. Genomic maps of long noncoding rna occupancy reveal principles of rna-chromatin interactions. Molecular cell, 44(4):667–678, 2011a. ISSN 10972765. URL http://www.sciencedirect.com/science/article/pii/S1097276511006800.

Ci Chu, Kun Qu, Franklin L Zhong, Steven E Artandi, and Howard Y Chang. Genomic maps of long noncoding rna occupancy reveal principles of rna-chromatin interactions. Molecular cell, 44 (4):667–678, 2011b.

G Felsenfeld, David R Davies, and Alexander Rich. Formation of a three-stranded polynucleotide molecule. Journal of the American Chemical Society, 79(8):2023–2024, 1957.

Phillip Grote, Lars Wittler, David Hendrix, Frederic Koch, et al. The tissue-specific lncrna fendrr is an essential regulator of heart and body wall development in the mouse. Developmental cell, 24(2):206–214, 2013. ISSN 15345807. URL http://www.sciencedirect.com/science/article/pii/S1534580712005862.

Mitchell Guttman and John L Rinn. Modular regulatory principles of large non-coding RNAs. Nature, 482(7385):339–46, February 2012. ISSN 1476-4687. doi: 10.1038/nature10887. URL http://www.ncbi.nlm.nih.gov/pubmed/22337053.

Sha He, Hai Zhang, Haihua Liu, and Hao Zhu. Longtarget: a tool to predict lncrna dna-binding motifs and binding sites via hoogsteen base-pairing analysis. Bioinformatics, 31(2):178–186, 2015.

Rory Johnson and Roderic Guig´o. The RIDL hypothesis: transposable elements as functional domains of long noncoding RNAs. RNA (New York, N.Y.), 20 (7):959–76, July 2014. ISSN 1469-9001. doi: 10.1261/rna.044560.114. URL http://www.pubmedcentral.nih.gov/articlerender.fcgi?artid=4114693&tool=pmcentrez&rendertyp

Tim R Mercer and John S Mattick. Structure and function of long noncoding rnas in epigenetic regulation. Nature structural & molecular biology, 20(3):300–307, 2013.

Tanmoy Mondal, Santhilal Subhash, Roshan Vaid, Stefan Enroth, Sireesha Uday, Bj¨orn Reinius, Sanhita Mitra, Arif Mohammed, Alva Rani James, Emily Hoberg, Aristidis Mous-takas, Ulf Gyllensten, Steven J M Jones, Claes M Gustafsson, Andrew H Sims, Fredrik Westerlund, Eduardo Gorab, and Chandrasekhar Kanduri. MEG3 long noncoding RNA regulates the TGF-β pathway genes through formation of RNA-DNA triplex structures. Nature communications, 6:7743, 2015. ISSN 2041-1723. doi: 10.1038/ncomms8743. URL http://www.nature.com/ncomms/2015/150724/ncomms8743/full/ncomms8743.html\nhttp://www.natur

Matthew D Simon, Charlotte I Wang, Peter V Kharchenko, Jason A West, Brad A Chapman, Artyom A Alekseyenko, Mark L Borowsky, Mitzi I Kuroda, and Robert E Kingston. The genomic binding sites of a noncoding rna. Proceedings of the National Academy of Sciences, 108(51): 20497–20502, 2011.

